# Detection of local growth patterns in longitudinally imaged low-grade gliomas

**DOI:** 10.1101/2022.04.24.488099

**Authors:** Chloe Gui, Jason Kai, Ali R. Khan, Jonathan C. Lau, Joseph F. Megyesi

**Author notes:** **Correspondence to:** Chloe Gui, MD and Jonathan C. Lau, MD, PhD, FRCSC Department of Clinical Neurological Sciences, Division of Neurosurgery, Western University, University Hospital, 339 Windermere Road, London, ON, Canada N6A 5A5. These authors were co-senior authors of this study: Joseph F. Megyesi and Jonathan C. Lau. **Funding:** C.G. received research scholarships from the Summer Research Training Program at the Schulich School of Medicine and Dentistry at Western University and the Mach-Gaensslen Foundation of Canada. J.C.L was funded through the Western University Clinical Investigator Program accredited by the Royal College of Physicians and Surgeons of Canada and a Canadian Institutes of Health Research Frederick Banting and Charles Best Canada Graduate Doctoral Award Scholarship. **Authorship:** C.G. and J.C.L. contributed equally to this work, designed and implemented the imaging pipeline, analyzed and interpreted the data, and wrote the manuscript. J.K. assisted with the pipeline and data interpretation. J.F.M. and A.K. interpreted the data and provided critical comments.

## Abstract

**Background:** Diffuse low-grade gliomas (LGGs) are primary brain tumors with infiltrative, anisotropic growth related to surrounding white and grey matter structures. In this study, we illustrate the use of deformation-based morphometry (DBM) as a simple and objective method to study the local change in growth patterns of LGGs.

**Methods:** An imaging pipeline was developed involving the creation of patient-specific average templates and nonlinear registration of pre-treatment follow-up MRIs to the average template. Jacobian maps were derived and analyzed to identify areas of tissue expansion and contraction over time.

**Results:** Our analysis demonstrates that tissue expansion occurs primarily around the edges of the tumor, while the lesion core and areas adjacent to obstacles, such as the skull, show no significant growth. Tumors also appeared to grow faster and predominantly in areas of white matter. Regions of the brain surrounding the lesion showed slight contraction over time, likely representing compression due to mass effect of the tumor.

**Conclusions:** We demonstrate that DBM is a useful clinical tool to understand the long-term clinical course of an individual’s tumor and identify areas of rapid growth, which can explain the clinical signs and symptoms, predict future symptoms, and guide targeted diagnostics and therapy.

**Highlights:**

1. Low-grade glioma expansion occurs primarily around the edges of the tumor.
2. Tumor cores and tissue next to obstacles show no significant growth over time.
3. DBM provides a clinically valuable assessment of local tumor growth and activity.

## Introduction

Diffuse low-grade gliomas (LGGs), classified as grade II gliomas by the World Health Organization, are primary brain tumors that arise from the glial cells of the brain.^1^ LGGs exhibit slow growth, expanding in diameter by 3.5 to 5 mm/year,^2–7^ but they eventually evolve into high grade tumors with poor outcomes. The disease often occurs in patients who are young and in relatively good health, typically presenting with seizures that can be medically controlled. Management of these patients is controversial due to the initial period of clinical stability despite inevitable transformation. With no randomized controlled trials, there is little concrete evidence that one particular management method is superior, though in recent decades, numerous retrospective studies have suggested that prompt, maximal, and even supratotal resection may prolong survival.^8–12^ On the other hand, some institutions, for certain cases, do watchful observation with regular imaging and follow-up, then intervene when the tumor is radiologically and/or clinically suspected to have progressed or transformed.^13–15^

Previous studies on the growth dynamics of LGGs have examined volume and diametric expansion on serial imaging. These studies have demonstrated that the tumors grow persistently, even if the patient appears clinically stable.^2–7^ Growth rate and size on index scans predict time to malignant transformation, clinical intervention, and overall survival, with slow-growing tumors leading to more favourable outcomes.^2–7^ While overall growth and volume is prognostically useful, these volumetric techniques capture only coarse changes and lack spatial information, failing to capture the anisotropic local growth of LGGs. Furthermore, tumor measurement by segmentation or by determining the mean diameter^16^ requires a trained individual. This process is time-consuming and subjective, leading to a degree of inter-rater and intra-rater variability.^17^

Deformation-based morphometry (DBM) is an automated, objective, and reproducible technique in the field of computational neuroanatomy.^18^ In the process of nonlinear registration, deformation fields are produced that describe the spatial transformation of voxels between scans, which characterize variations in anatomy. This method allows the quantification of neuroanatomical variation at the voxel level across the entire brain without bias for any particular structures.^18^

Longitudinal imaging studies have used DBM to quantify structural changes in the brain over time and in various neurological and psychiatric diseases in both animal^19^ and human populations.^18,20–22^ Studies on tumors with DBM are less common, but the method has been applied to assess breast cancer response to chemotherapy^23^ and esophageal cancer response to chemoradiotherapy.^24^ There have been no studies to date using DBM to examine the growth of LGGs. Due to their gradual growth, anisotropic invasion patterns and relatively homogenous hyperintensity on T2-weighted imaging, we anticipated this technique would be particularly suitable for morphometric measurements of these tumors. In this study, we applied DBM to quantify the morphological changes of LGGs from a previously-described cohort that was serially-imaged during a period of watchful observation period prior to oncological treatment.^15^

## Methods

### Patient Population

We previously described the natural history and tumor growth trajectories on serial MRI of LGG patients that were managed by watchful observation over a median of 79.7 months.^15^ From a database of 842 brain tumor patients that were under the care of senior author J.F.M. from April 2004 to July 2017, 74 had LGGs. Of these 74 patients, patients with at least 8 MRIs before oncological treatment (including chemotherapy, radiation, surgical resection) were selected for further analysis of tumor growth over time for a total of 10 LGG patients. T2-weighted MRIs of these patients were retrieved from the Picture Archive and Communications System, and clinical data was obtained from electronic medical records. The in-house DBM analysis pipeline we developed was applied to each patient. All patients were at least 18 years old, and all tumors were histopathologically confirmed to be World Health Organization grade II gliomas. This study was approved by the Western University and London Health Sciences Centre Research Ethics Board.

### MRI Acquisition & Tumor Segmentation

MRI studies were acquired every 3-12 months using a clinical 1.5 T GE Twin or 1.5 T GE CVMR scanners (GE Healthcare, Milwaukee, Wisconsin, USA). To create tumor masks that aid in registration, the tumors on MRI were manually segmented using ITK-SNAP (version 3.6.0).^25^ Inter-and intrarater Dice similarity coefficients were calculated using the Convert3D Medical Imaging Processing Tool, developed by ITK-SNAP (www.itksnap.org), to quantify reproducibility. Tumor masks were used in all rigid registration steps and to define the tumor region during analysis. Eroded tumor masks were used to identify the core versus peripheral regions. To define tumor areas adjacent to and away from the skull, whole-brain masks were slightly eroded, and areas of overlap between the eroded brain mask and tumor mask were defined as areas away from the skull.

### Patient Template Creation

An average template scan was created for each patient using all of the patient’s follow-up scans prior to oncological treatment. Each scan was first corrected for intensity nonuniformity using the N4 algorithm.^26^ Scans were then rigidly registered to the MNI ICBM 2009c Nonlinear Asymmetric 1×1×1mm T2-weighted template^27,28^ using the Advanced Normalization Tools (ANTs) package,^29^ specifically the antsRegistration tool, to resample the resolution of all scans to 1×1×1 mm. Next, the middle follow-up session scan for each patient was chosen as the reference image, and each scan was rigidly registered to the patient’s middle scan to bring all scans to the same patient MRI space. Registered scans were visually inspected for acceptable registration accuracy. Scans were then transformed using the antsApplyTransform tool and affine matrices produced during the previous registration steps to bring the scans into their respective patient space.^29^ These images were then used to create the patient-specific average template with the ANTs Multivariate Template Construction tool.^29^ A deformable registration step was performed to register follow-up scans to their respective average template, resulting in a deformation field describing the global and local mapping between each individual scan and the average template for the patient.^29,30^

### Jacobian Determinants

The Jacobian determinants provide a measure of local contraction or expansion in a voxelwise manner. They are partial derivatives of the deformation field and were computed from the registration of follow-up scans to their respective patient-specific average template. To obtain a normal distribution, the Jacobian determinants were log-transformed before analysis. The log Jacobian is also simpler than the Jacobian to interpret, with a value greater than zero indicating local tissue expansion and a value less than zero indicating local tissue contraction. Values equal to zero correspond to no local volume change. For comparative analysis of local growth, tumors masks were eroded to identify the peripheral and cores regions of the lesion. Whole-brain and eroded brain masks were used to isolate the regions of the tumor adjacent and away from the skull.

### Statistical Analysis

For each patient, linear models were fit at each voxel to analyze change in the Jacobian over time. Here, we also describe the coefficient (estimate or slope) of the linear regression as a measure of time-related change. Statistical results were corrected for multiple comparisons using the false discovery rate procedure.^31^ Analysis was performed in R (version 3.5.1).^32^ The fslr package was used to handle NifTI files during the analyses;^33,34^ linear modeling was performed with the native Stats package, and the ggplot2 package was used for graphing.^35^ The Wilcoxon test was used to compare mean Jacobian change over time between different tumor regions.

## Results

### General Growth Patterns

Patient demographics and tumor characteristics are presented in Table 1, and clinical patterns of tumor growth have been previously discussed.^15^ Axial images of each subject with an overlay of longitudinal voxelwise change in the tumor region are shown in Figure 1 to illustrate general growth patterns of the lesions. Jacobians significantly increased with time primarily at the peripheral regions of the tumor, indicating expansion relative to the patient average template (Fig. 1). Local changes in these peripheral regions were significantly greater than at the core of the tumor (Fig. 2,3B). Furthermore, areas of the tumor adjacent to the skull had significantly smaller local changes over time compared to areas of the tumor away from the skull (Fig. 1,2,3C). A similar of slower growth was seen in tumor regions next to the falx (Fig. 1). Areas of relative longitudinal contraction were notable among brain tissue just beyond the borders of the lesion, suggesting compression from the mass effect of the tumor (Fig. 2,3E).

**Table 1.** Patient demographics, tumor characteristics, and imaging details

| Patient | Age | Sex | Tumor Location | VDE (mm/year) | MRIs (n) | Follow-Up (months) |
| --- | --- | --- | --- | --- | --- | --- |
| 1 | 36 | M | Left parietal | 2.20 | 18 | 97.9 |
| 2 | 43 | F | Right frontal-parietal parasagittal | 1.82 | 11 | 108.3 |
| 3 | 63 | M | Left frontal-temporal (insular) | 4.29 | 11 | 90.8 |
| 4 | 22 | M | Right parietal | 1.95 | 12 | 97.6 |
| 5 | 37 | F | Right parietal | 2.99 | 8 | 41.2 |
| 6 | 42 | F | Left temporal occipital | 1.56 | 14 | 113.8 |
| 7 | 54 | M | Right frontal parasagittal | 5.02 | 13 | 49.8 |
| 8 | 39 | M | Right frontal-temporal (insular) | 2.14 | 8 | 63.5 |
| 9 | 53 | M | Right frontal | 3.31 | 12 | 58.6 |
| 10 | 45 | M | Left frontal | 1.97 | 11 | 39.8 |

VDE: velocity of diametric expansion

**Figure 1.**
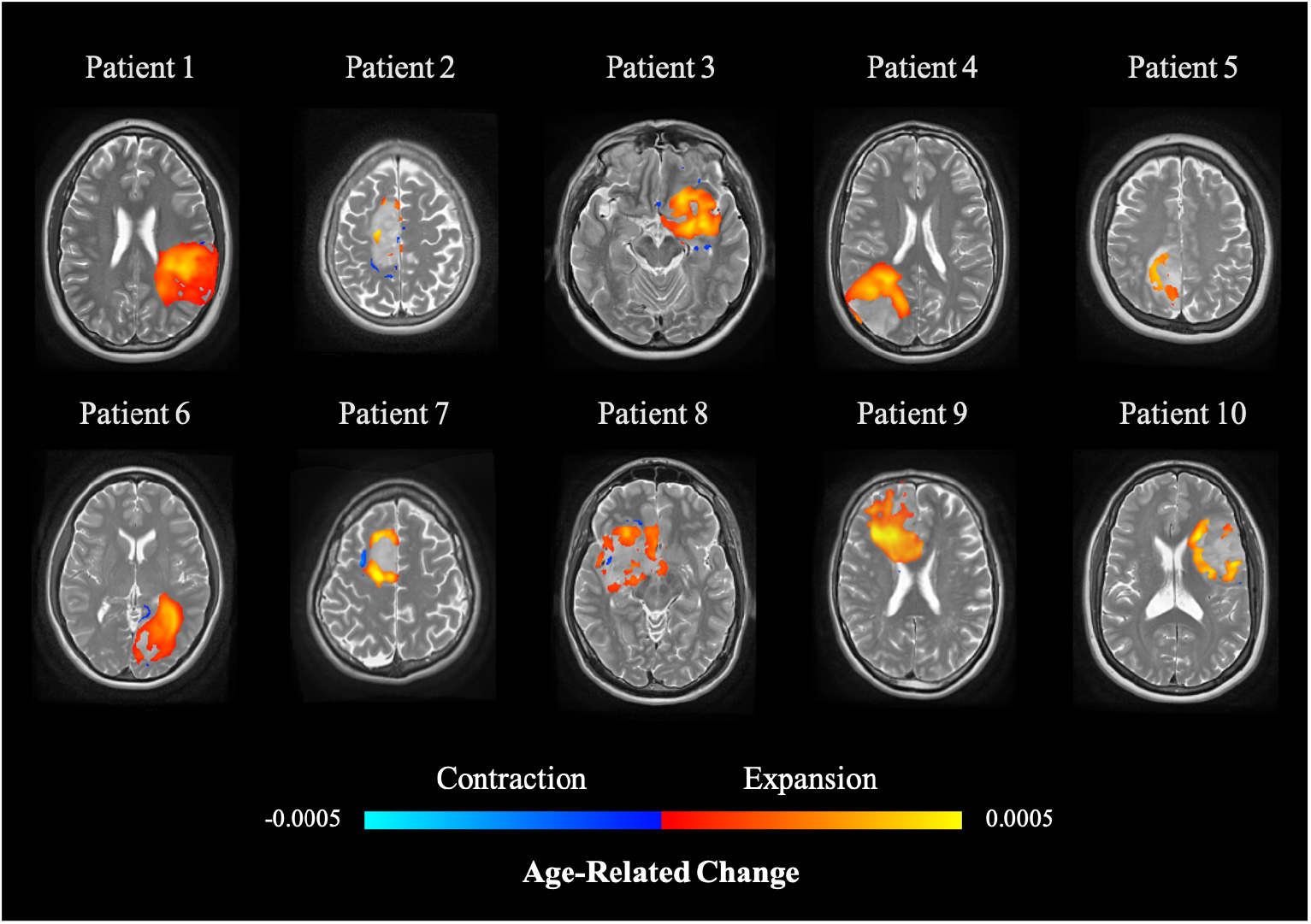
Axial MRI images of the centre of each subject’s tumor are displayed. The linear model coefficient of Jacobian over time at each voxel is overlaid, representing areas of significant morphometry change. Areas of greatest change are primarily at the perimeter of the lesions, and some slight contraction is visible just adjacent to the tumor. Expanding voxels are represented in red to yellow, and contracting voxels are represented in blue.

**Figure 2.**
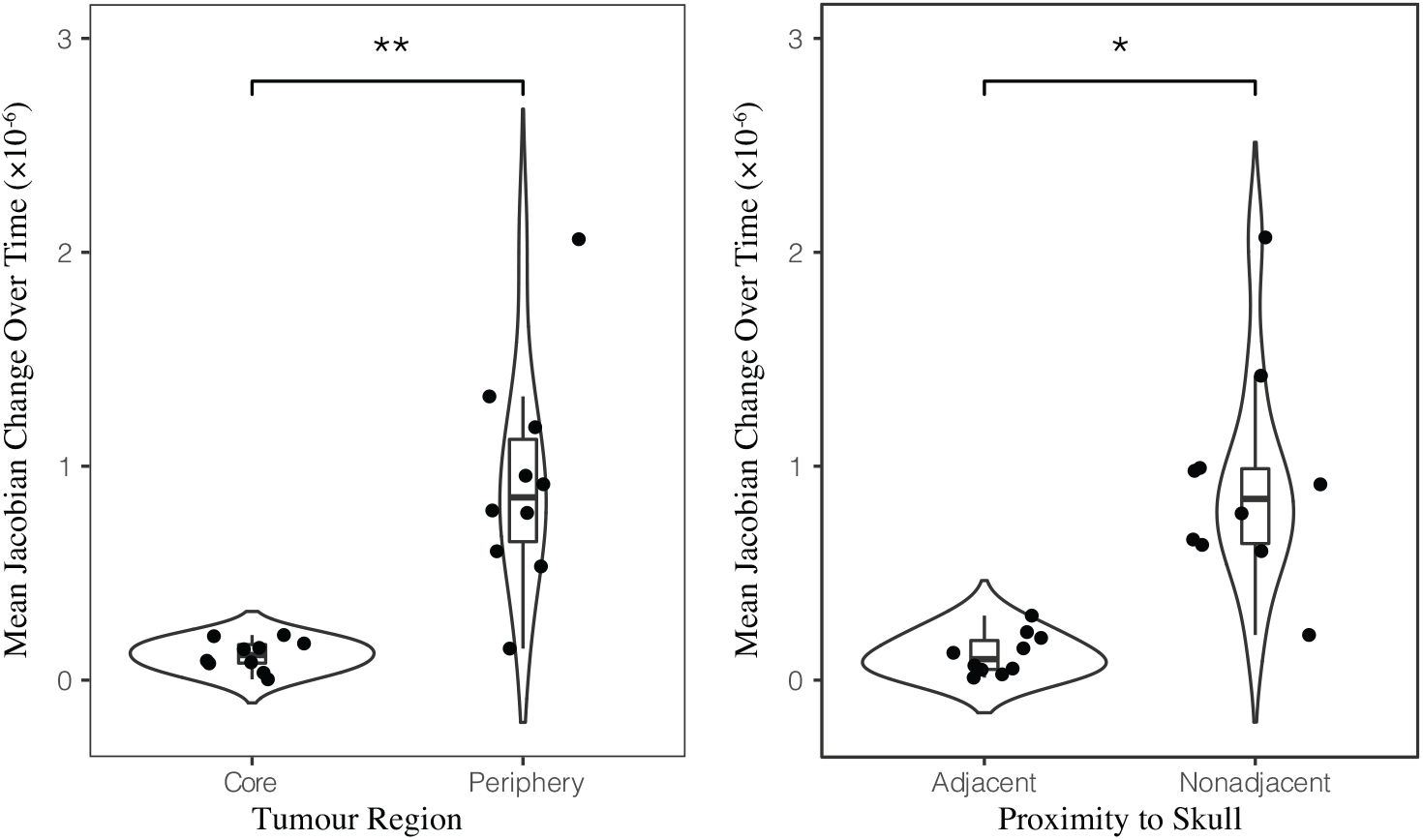
(A) Voxelwise expansion in Jacobian over time is significantly greater at the periphery of the tumor region compare to the tumor core. (B) Tumor expansion is significantly more rapid in areas away from the skull compare to areas adjacent to the skull. Black bars indicate the mean Jacobian change over time among each group. (* *p* < 0.001, \**p* = 0.015).

**Figure 3.**
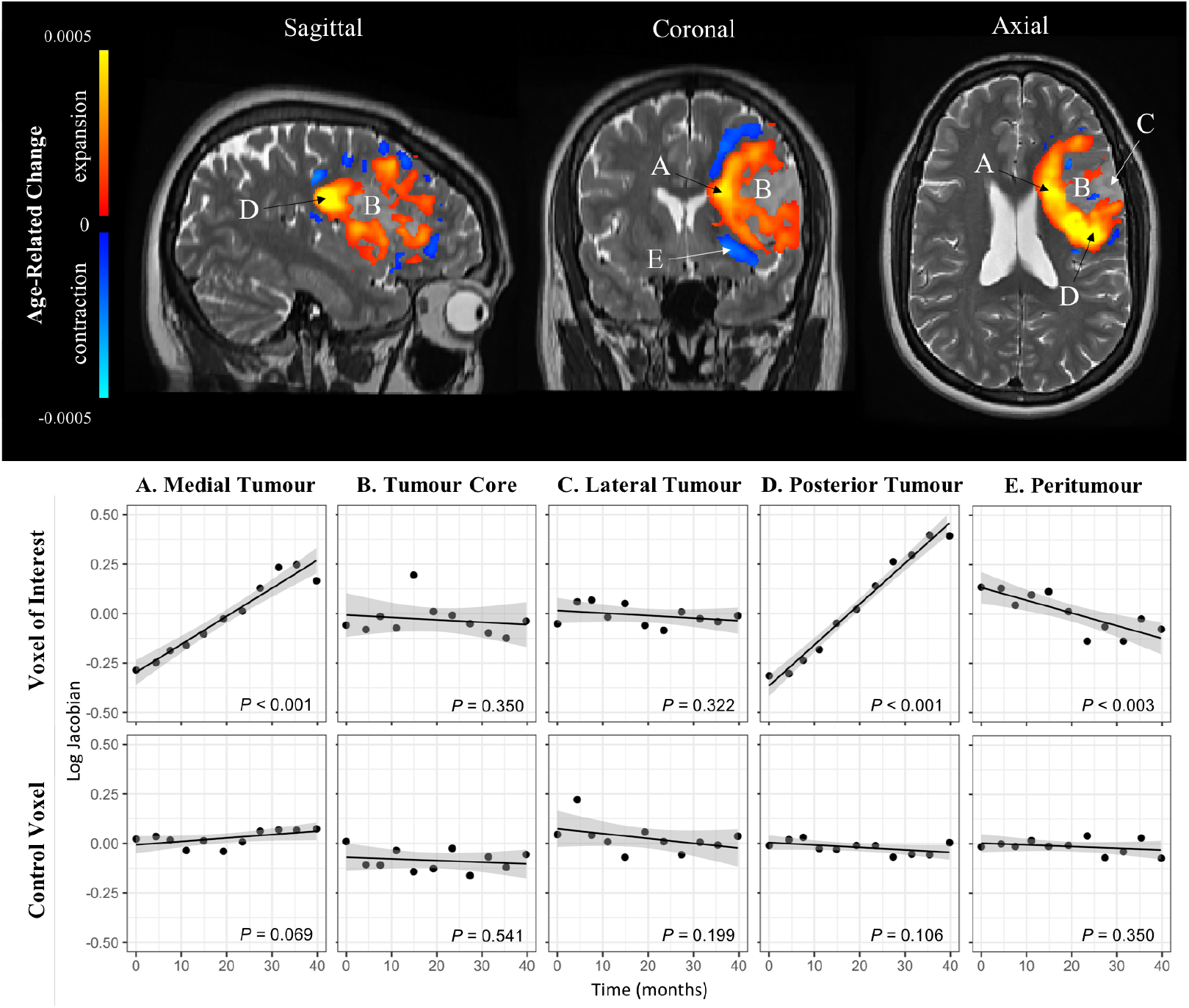
MRI slices of a representative subject (Patient 10 of Figure 1) and overlay of the linear modeling map of Jacobian change over time, with areas of red and yellow representing expansion (log Jacobian > 0) and areas of blue representing contraction (log Jacobian < 0). Linear modeling of voxelwise log Jacobian over time demonstrates volume expansions in the (A) medial and (D) posterior tumor and stability in the (B) tumor core and (C) lateral tumor. (E) indicates an area of brain tissue contraction due to presumed mass effect from the tumor. Areas of interest were compared to contralateral control voxels. Linear regression did not reveal any significance at any of the control voxels.

### Analysis of Representative Case

Figure 3 highlights several regions of interest in a representative patient (Patient 10). In the region of interest, 16.1% of voxels had Jacobians that significantly changed over time. Of these, 95.45% of the voxels demonstrated expansion over time, and 4.55% of the voxels indicating tissue compression. Clusters of voxels with local expansion were notable along the medial (Fig. 3A) and posterior aspects (Fig. 3D) of the tumor, whereas the lateral edge of the tumor adjacent to the skull showed no significant change (Fig. 3C). The core region of the tumor did not demonstrate any volumetric change, suggesting stability over time (Fig. 3B). The voxels of interest were compared to control locations on the contralateral hemisphere, where there were no significant morphological changes over time. Our analysis also yielded a few regions of significant tissue contraction around the tumor (Fig. 2E). These areas were outside the visible boundaries of the lesion, suggestive of mass effect on brain regions around the tumor. On an axial view, they were notable anterior and posterior to the tumor, and on a coronal view, they were predominantly dorsomedial and ventromedial to the tumor.

## Discussion

In this study, we demonstrated the feasibility of DBM analysis on tumor growth in patients with LGGs and found that linear regression of Jacobians over time reveals sites of local tumor expansion alongside areas of relative stability and peritumor contraction of brain tissue.

### Tumor growth

Local areas of significant expansion were found in all tumors (Fig. 1,3), primarily near the edges of the tumor and away from impeding structures such as the skull (Fig. 2). The tumor core was generally stable over time, and some patients demonstrated compression of brain parenchyma outside of the tumor borders, likely representing the compression of normal brain tissue by the enlarging tumor.

Two previous studies sought to quantify tumor change during chemotherapy and radiation treatment.^23,24^ One study found that deformable registration to be a useful tool in recognizing breast tumour change in longitudinal MRIs.^23^ Another group analyzed advanced esophageal tumors between CT scans and noted that the mean and minimum Jacobian determinants were important parameters in differentiating responders from non-responders.^24^ Future studies on LGG shrinkage during chemoradiation could further clarify whether the Jacobian could be a useful volume change parameter in the context of treatment.

### Clinical application of deformation-based morphometry

Imaging follow-up is an important aspect of LGG management. For neuro-oncology teams that choose to observe LGG patients during the clinically stable phase of the disease, significant changes on imaging may be an indication to treat, and patients are followed with imaging post-operatively as well. Lesions are not typically consistently measured on imaging, and about 20% volume change is necessary for visual appreciation of growth.^36^ DBM is highly sensitive and quantitative, allowing a more accurate method of assessing growth, and does not require the labour-intensive segmentation or manual diameter measurements that current research studies use.^2–7^ The clear differentiation between areas of rapid versus subtle growth can inform physicians of the areas of clinical concern, which may lead to future symptoms or explain a patient’s current presentation. The areas of high growth are also likely the most aggressive parts of the tumor and may be ideal locations for biopsy targeting to avoid under grading. Furthermore, awareness of which local areas of the tumor are slow-growing may inform clinical management, as physicians may feel more comfortable observing patients rather than treating promptly if important or eloquent structures are not in immediate danger of invasion. Given the risks of brain surgery, DBM provides an informative and clinically valuable assessment of tumor growth and activity.

### Limitations

Several limitations have been discussed in our previous reporting of this cohort of patients. In brief, the group consists of LGGs that are slower-growing, and the specific growth patterns of the tumors presented in this study may not be generalizable to all LGGs.^15^

Limitations surrounding our pipeline and analysis are as follows. The registration portion of the pipeline used tumor masks that were manually segmented to improve registration and define the region of interest. Segmentation inaccuracies contribute to a certain degree of error that we have previously quantified with the Dice similarity coefficient (interrater: 0.867, intrarater: 0.914).^15^ Registration inaccuracies may also contribute to errors in the deformation maps; every scan was visually assessed post-registration by experts to minimize this.

## Conclusions

Understanding tumor growth dynamics could lead to improvements in the management and treatment of LGGs. Here, we observed the anisotropic growth pattern of LGGs and demonstrated how DBM highlights regions of the tumor that are rapidly growing. We found that the areas of most rapid expansion were in the peripheral regions of the lesion, areas that were away from obstacles such as the skull, and areas that colocalized with white matter tracts. We also found that overall Jacobian were not good indicators of growth for this subset of tumors and are more valuable for describing local changes. Overall, DBM is a simple, objective tool with the potential to provide clinically valuable information for LGG researchers and clinicians.

## Acknowledgements

This work was previously presented as a platform presentation at the 2019 Canadian Neurological Sciences Federation in Halifax, Nova Scotia, Canada.

