## Supplementary Figure 1 for "Detection of local growth patterns in longitudinally imaged low-grade gliomas"

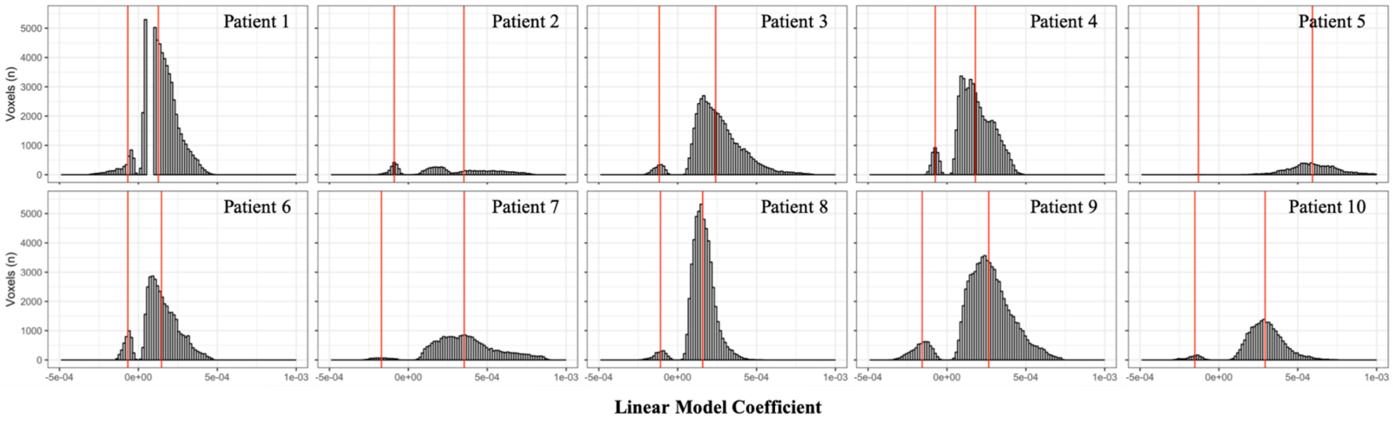


**Supplementary Figure 1.** Histograms depicting the distribution of linear model coefficients in the region of interest around the tumor all subjects. Red vertical lines indicate the mean among positive coefficients and negative coefficients. Most subjects had a large proportion of positive coefficients, representing voxels that are expanding over time. A smaller peak of negative coefficients can be seen in most subjects, representing areas of tissue contraction over time. The magnitude of the coefficient corresponds to the slope of the linear model, or how rapid expansion or contract at a given voxel is.
